# Individual variability in functional connectivity architecture of the mouse brain

**DOI:** 10.1101/2020.03.16.993709

**Authors:** Eyal Bergmann, Xenia Gofman, Alexandra Kavushansky, Itamar Kahn

## Abstract

The functional organization of brain networks can be estimated using fMRI by examining the coherence of spontaneous fluctuations in the fMRI signal, a method known as resting-state functional connectivity MRI. Previous studies in humans reported that such functional networks are dominated by stable group and individual factors, demonstrating that fMRI is suited to measuring subject-specific characteristics, and suggesting the utility of such precision fMRI approach in personalized medicine. However, mechanistic investigations to the sources of individual variability in health and disease are limited in humans and thus require animal models. Here, we used repeated-measurement resting-state fMRI in awake mice to quantify the contribution of individual variation to the functional architecture of the mouse cortex. Comparing the organization of functional networks across the group, we found dominant common organizational principles. The data also revealed stable individual features, which create a unique fingerprint that allow identification of individual mice from the group. Examining the distribution of individual variation across the mouse cortex, we found it is homogeneously distributed in both sensory and association networks. Finally, connectome-based predictive modeling of motor behavior in the rotarod task revealed that individual variation in functional connectivity explained behavioral variability. Collectively, these results show that mouse functional networks are characterized by individual variations suggesting that individual variation characterizes the mammalian cortex in general, and not only the primate cortex. These findings lay the foundation for future mechanistic investigations of individual brain organization and pre-clinical studies of brain disorders in the context of personalized medicine.

## Introduction

A fundamental question in brain research is what makes individuals different from each other. This question can be addressed at different levels of organization, starting from genetics or neurotransmitters, going through structural or functional measures of brain regions and networks, and ending at behavioral phenotypes or clinical outcomes. In humans, a common approach to study brain organization in individuals is resting-state functional connectivity fMRI (fcMRI), which estimates functional connectivity between regions based on coherent spontaneous fluctuation in the fMRI signal^1–3^.

Previous human fcMRI studies demonstrated that this measure is stable over time and can be used to characterize individual differences^4–8^. These works revealed that such differences are spread heterogeneously across the human cortex, demonstrating increased variability in association networks, and correlation with distal connectivity and cortices that underwent expansion and elaboration relative to non-human primates and lower mammals. Moreover, individual variations in functional connectivity were shown to predict individual activity patterns in task conditions^9,10^ and behavioral performance^11–13^. Finally, recent studies in patients with neuropsychiatric disorders reported that individual functional connectivity patterns can be used as a biomarker for diagnosis and treatment optimization^14–16^, key features of personalized medicine.

While individual differences in functional connectivity were thoroughly characterized in humans, including identification of sources of intra-subject variability^17^, a dissection of mechanisms relies on animal models. Such investigation demands adequate sample size that is hard to achieve in studies in non-human primates and may involve genetic manipulations and molecular techniques that are more readily accessible in rodent models, and specifically in mice. Previous fcMRI studies in anesthetized mice demonstrated reproducible resting-state networks^18,19^, applications to mouse models of neuropsychiatric diseases^20–22^, and correlations between functional connectivity and behavioral measures^23–26^. However, characterization of individual differences in functional connectivity is based on repeated data acquisition that can control for measurement instability. Since this experimental design is hard to achieve in anesthetized animals, such studies are more suitable to awake mouse imaging. We have previously established fcMRI experiments in awake head-fixed mice^27,28^ and used repeated-measurement designs to link individual differences in structural and functional connectivity^29^. However, a detailed analysis of individual variability of functional connectivity in the mouse brain and its relevance to behavior has heretofore not been demonstrated.

Here we used repeated-measurement resting-state fMRI to characterize individual variation in functional connectivity in the mouse cortex. We show that despite the mouse cortex reduced complexity relative to the human homologue and the animals being genetically identical, it is also characterized by individual variation, allowing identification of specific mice from a group. Then, we characterize factors affecting identification accuracy, and examine the distribution individual variability in sensory and association networks. Finally, we link individual differences in functional connectivity to behavioral variability in the rotarod task. Collectively, these findings indicate that mouse functional networks are characterized by behaviorally relevant individual variation and lay the foundation for mechanistic investigations of sources of individual variability and pre-clinical studies of brain disorders in the context of personalized medicine.

## Results

### Connectivity-based identification of individual mice

Data of the study consisted of nineteen F1 C6/129P (male, age 9–12 weeks), which underwent multiple fcMRI sessions during passive wakefulness as previously described^29^, followed by behavioral testing in the rotarod task^30^. After exclusion of sessions with image artifacts or excessive motion (see methods), the final dataset included 16 mice with six sessions (each comprising ~30 minutes of data), which were split to two halves of three sessions each to examine the group and individual similarities in the mouse functional connectome. For the comparison between functional connectivity and behavior, two additional mice with 4–5 sessions were included. In this analysis, data from all sessions were averaged, resulting in a single connectivity matrix per mouse.

Functional connectivity matrices were built based on the Allen Common Coordinate Framework reference atlas (CCFv3) downsampled to fMRI resolution (**Fig. 1A**), including 43 cortical parcels per hemisphere, which were divided to six modules (Prefrontal, Lateral, Somatomotor, Visual, Medial and Auditory) based on anatomical connectivity patterns^31^. The two connectivity matrices of each mouse were compared to each other and to all other connectivity matrices of the other mice (representative matrices are presented in **Fig. 1B**) by calculating the Fisher-transformed correlations between all edges in the connectome. The resulting network similarity matrix (**Fig. 1C**) diagonal represents the similarity within individual, while rows and columns represent similarity between specific mouse and all other mice in the group. This matrix was used for quantification of individual variation in the connectome by comparing group and individual network similarities^7^. We found substantial group similarity (mean z(r) = 0.9), which indicates mouse connectomes share a common structure. Nevertheless, we also found that individual similarity (mean z(r) = 1.08) is significantly higher than group similarity (two-tailed paired student *t*-test: t_(15)_ = 7.32, P < 0.001, Cohen’s *d* = 1.83), demonstrating larger values in all mice (**Fig. 1D**), and indicating that fcMRI can capture individual variability in the mouse connectome.

**Figure 1.**
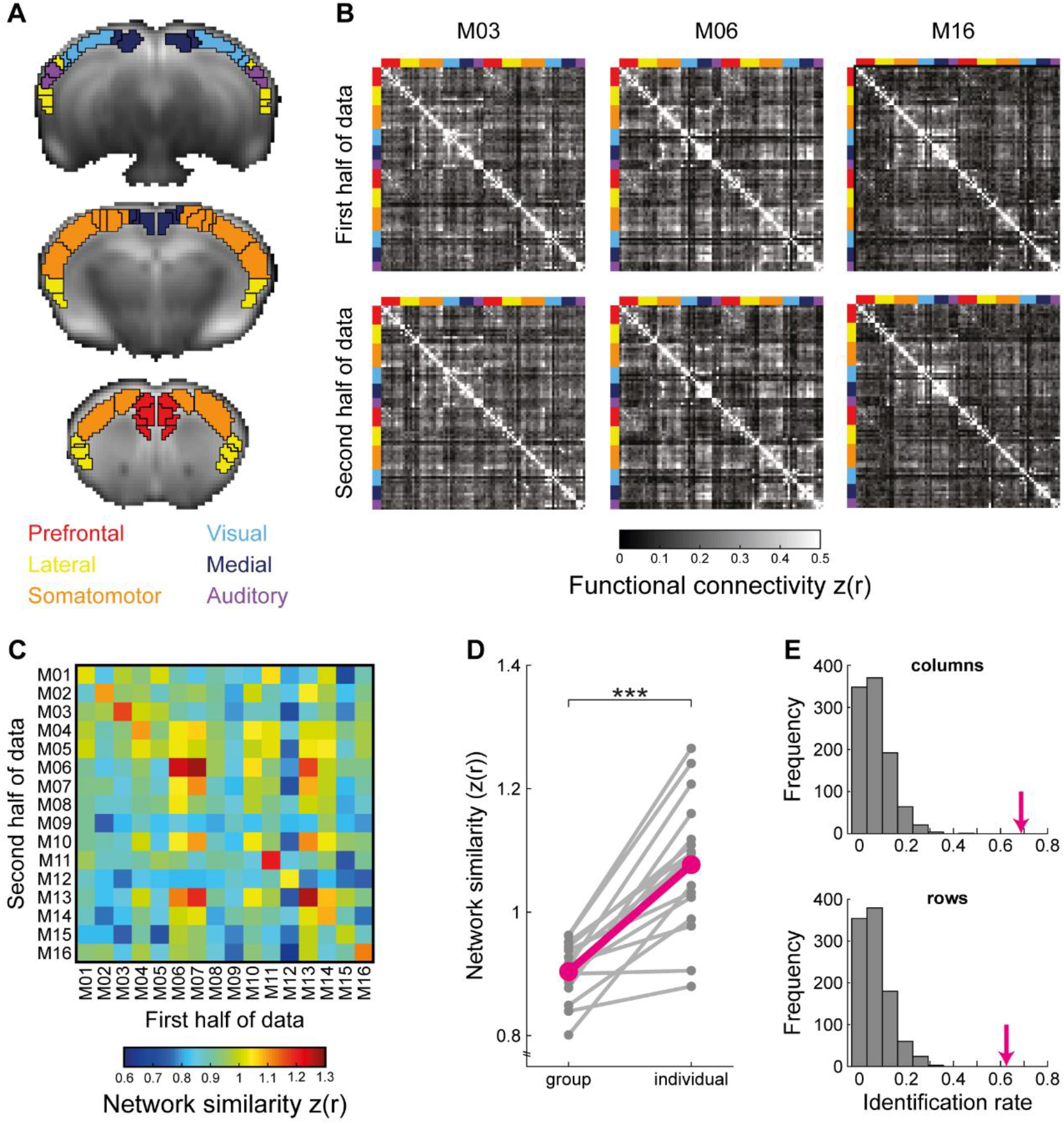
Functional connectome fingerprinting in the mouse cortex. (A) Anatomical parcellation of the mouse cortex overlaid on average fMRI image at native resolution. Eighty six cortical labels (black borders) were taken from the Allen Mouse Brain Atlas CCFv3 and divided to six modules (label color) based on their anatomical connectivity profiles^31^. (B) Individual functional connectomes of three representative mice from the cohort demonstrate unique patterns that are consistent between the first and second halves of data. The identity of the anatomical modules of each node is designated by the color. (C) A network similarity matrix in which each cell represents the similarity between two connectomes. The matrix is asymmetric as columns represent the similarity between the first half of data of each mouse and the second half of the data of all the other mice, while rows represent the similarity between the second half of data of each mouse and the first half of data of all the other mice. The values along the diagonal represent individual similarity, which is the correlation between the edges of the two connectivity matrices of the same mouse. (D) A comparison between group and individual similarities demonstrate higher similarity between connectomes from the same mouse (***P < 0.001), the *magenta* line depicts the average group and individual network similarities. (E) True connectome-based identification rates *(magenta)* were compared to 1000 iterations of identity shuffling indicating that the mouse functional connectome acts as a unique fingerprint.

The significant difference between group and individual network similarities means that on average, connectivity matrices from the same mouse are more similar than connectivity matrices from different mice. A more stringent criterion for connectome-based fingerprinting is successful identification of specific mouse from a group^12^. Namely, that the similarity between the two connectivity matrices of the same mouse must be higher than the similarity to all other connectivity matrices in the group. To test the feasibility of functional connectome-based identification in mice, we calculated the fraction of mice in which the values along the diagonal of the similarity matrix were higher than other values in each row and column. We found that the rate of successful identification of mice in the first and second halves of data was 68.75% and 62.5% out of the 16 mice, respectively (**Fig. 1E**). To formally test the significance of those rates, we used a shuffling procedure, calculated the distribution of identification rates when assigning random identities in either the first or second halves of connectivity matrices, and found that the observed identification rates are significantly higher than values in all 1000 shuffled iterations (P < 0.001, **Fig. 1F**). This result indicates that that the mouse functional connectome acts as a unique fingerprint.

### Factors contributing to characterization of individual variation

Studies in humans demonstrated that successful identification^12^ and precise characterization of individual variation^7^ depend on the amount of fcMRI data available per participant. Therefore, leveraging the repeated-measurement design of our fcMRI experiment, we sought to characterize how much data is needed to stably characterize the functional cortical organization in individual mice.

In our original analysis (**Fig. 1**) two average connectivity matrices were calculated per mouse by splitting its six sessions to two halves, controlling for the total number of included frames per half (see Methods). In the current analysis we built a set of connectivity matrices using all combinations of one (n = 15), two (n=45) or three sessions (n=20) per mouse, and examined network similarity and identification rates as a function of the number of sessions averaged per connectome (**Fig. 2a)**. First, we calculated group and individual network similarity values for different number of included sessions and submitted the results to a repeated-measures ANOVA (corrected with the Huynh-Feldt method) with Individuality and Number of Sessions as within mouse factors. We found significant effects of both factors (Individuality: *F*_(1, 15)_ = 34.28 P < 0.001, ε_H-F_ = 1, η^2^ = 0.71; Number of Sessions: *F*_(2, 30)_ = 7318.76, P < 0.001, ε_H-F_ = 0.51, η^2^ = 0.998), confirming that network similarity between connectivity matrices from the same mouse is higher than group similarity, and that increasing the amount of data per mouse improves network similarity estimation. Importantly, we also found a significant interaction between Individuality and Number of Sessions (*F*_(2, 30)_ = 109.87, P < 0.001, *ε*_H-F_ = 0.5, η^2^ = 0.887), indicating that increasing the amount of data per mouse preferentially increases individual over group network similarity values. In agreement with this finding, the results of the identification analysis revealed that increasing the number of sessions improve identification (two-tailed unpaired student *t*-test: two sessions vs. one session: t_(58)_ = 9.08, P < 0.001, Cohen’s *d* = 2.9; three sessions vs. one session: t_(33)_ = 15.53, P < 0.001, Cohen’s *d* = 5.3; three sessions vs. two sessions: t_(63)_ = 8, P < 0.001, Cohen’s *d* = 2.15). Collectively, these analyses indicate that repeat-measurement fcMRI designs supported by awake imaging are useful for characterizing individual functional connectivity patterns.

**Figure 2.**
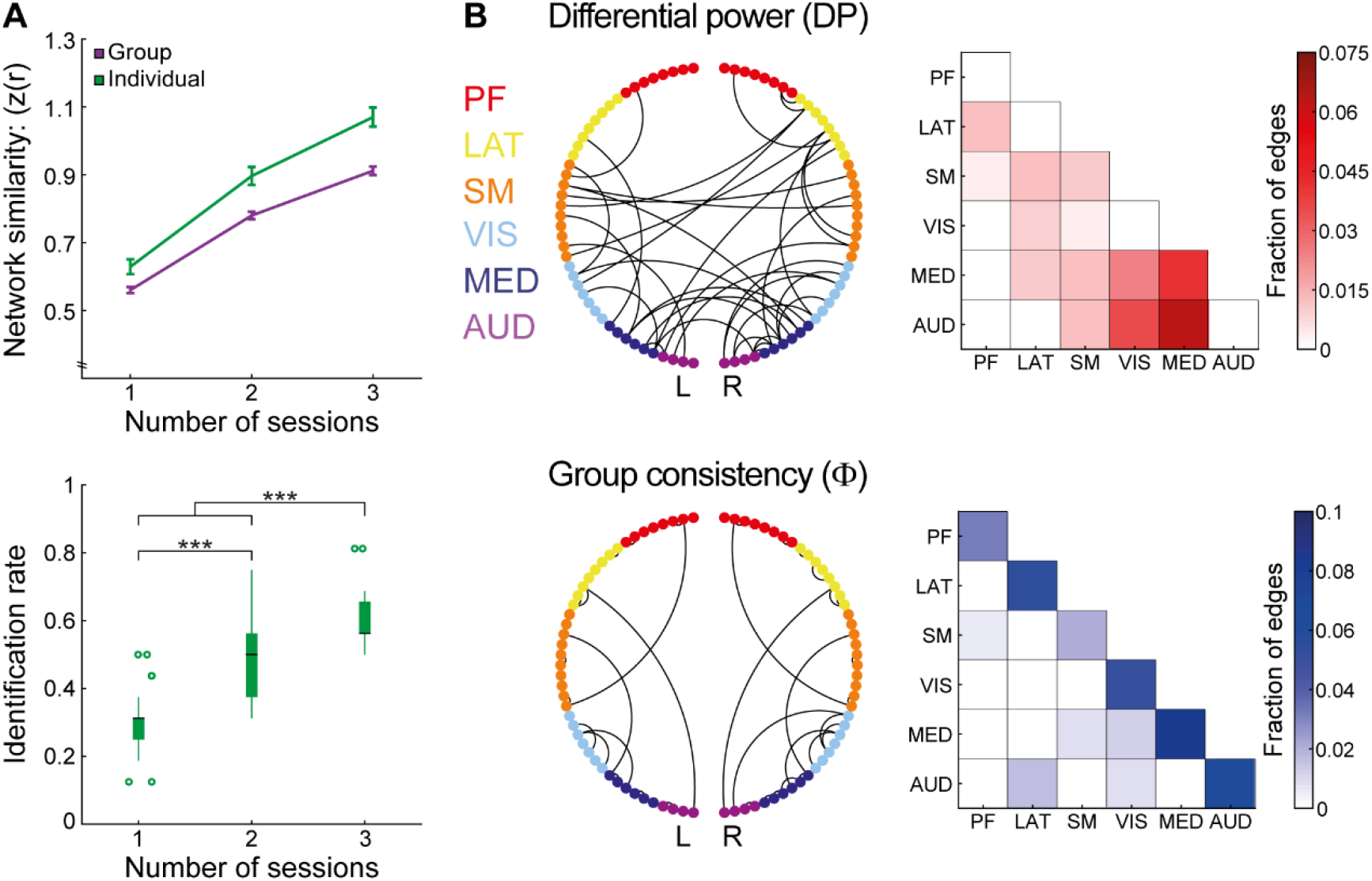
Factors contributing to characterization of individual variation. (A) Network similarity values *(top)* and identification rates *(bottom)* as a function of number of included sessions in the two connectivity matrices of each mouse. All combinations of data sampling of one, two or three sessions were calculated and compared using repeated-measures ANOVA (network similarity, see text) or two-tailed unpaired student *t-* test (identification rate, *** P < 0.001). error bars *(top)* represent the standard error of the mean; boxplots *(bottom)* represent the median (center line), interquartile range (box limits); 1.5 × interquartile range (whiskers) and outlier (points). (B) Unique (DP, *top*) and consistent (Φ, *bottom)* edges in individual connctomes. For visualization purposes, both measures were thresholded at the 99^th^ percentile. In the circle plot *(left)* the 86 nodes are organized based on their anatomical module identity and cortical hemisphere; lines indicate edges. In the matrices *(right),* the fraction of edges connecting between and within modules is color coded with darkly shaded cells representing higher DP (*top*) and Φ *(bottom)* values. PF, prefrontal; Lat, lateral; SM, somatomotor; VIS, visual; MED, medial; AUD, auditory.

After establishing that individual functional connectivity profiles act as a unique fingerprint in mice, we sought to characterize which brain connections contribute to individual variation. Therefore, we replicated that analysis of Finn et al.^12^ and characterized which edges possess high differential power (DP) that contribute to identification, and which edges possess high group consistency (Φ) within a mouse and across the group. We derived those values for all edges in the connectivity matrix, and determined which edges were in the top 1% of each measure (**Fig. 2b**). We found that most edges in the top 1% of DP are related to the posterior modules, namely, Visual, Medial and Auditory, and include many inter-hemispheric and inter-module connections. In contrast, edges in the top 1% of Φ came from all modules and included mainly intra-hemispheric intra-module connections. In the original analysis of Finn et al.^12^, high DP were observed in high-order fronto-parietal connections, while high Φ were observed in inter-hemispheric connectivity within the Somatomotor and Visual networks. Comparing the analyses across species, we conclude that while the mouse data recapitulate the Φ difference between intra- and inter-module connections, and show DP bias towards specific modules, it seems to not show a difference between sensory and association regions. Therefore, we sought to directly examine individual variation in those qualitatively different brain systems.

### Individual variation across brain systems

To better characterize individual variation in sensory and association systems in the mouse cortex, we sought to examine whether functional connectivity profiles within these systems differ in their identifiability or group and individual network similarities. Therefore, we defined each cortical node as either sensory (Somatomotor, Visual, and Auditory modules) or association (Prefrontal, Lateral, and Medial modules) and derived two network similarity matrices (**Fig. 3A**). Then, we calculated group and individual network similarities for each system (**Fig. 3B**), and found higher values compare to the original full connectivity matrix (all connections) in both group (two-tailed paired student *t*-test: all connections vs. association: t_(15)_ = 26.9, P < 0.001, Cohen’s *d* = 6.73; all connections vs. sensory: t_(15)_ = 18.44, P < 0.001, Cohen’s *d* = 4.61) and individual similarities (all connections vs. association: t_(15)_ = 8.92, P < 0.001, Cohen’s *d* = 2.23; all connections vs. sensory: t_(15)_ = 11.59, P < 0.001, Cohen’s *d* = 2.9). Comparison between association and sensory systems reveal significant difference in group network similarity (t_(15)_ = 4.96, P < 0.001, Cohen’s *d* = 1.24), but no difference in individual network similarity (t_(15)_ = 0.56 P, = 0.583, Cohen’s *d* = 0.14; all values were corrected for multiple comparison using false-discovery rate based on the Benjamini-Hochberg method). Examining the normalized relative effect magnitude of individuality (**Fig. 3C**), we found no difference between sensory and association systems (t_(15)_ = 1.803, P = 0.137, Cohen’s *d* = 0.45). Nevertheless, comparison to the full connectivity matrix revealed lower effect magnitude of individuality in association (t_(15)_ = 3.02, P = 0.026, Cohen’s *d* = 0.75), but not sensory (t_(15)_ = 0.134, P = 0.85, Cohen’s *d* = 0.03), systems. Finally, we carried out the identification analysis for each type of connectome and found similar identification rates in sensory and association systems (**Fig. 3D**), which were only slightly lower than the original values yielded by the full connectivity matrix. Collectively, these analyses indicate that connections between sensory and association regions, which reflect inter-module connectivity, are less consistent across the group. Importantly, the results indicate that the relative effect magnitude of individuality on functional organization of the mouse cortex is more modest compared to humans^7^, and the qualitative differences between sensory and association networks in the human brain are not well recapitulated in mice.

**Figure 3.**
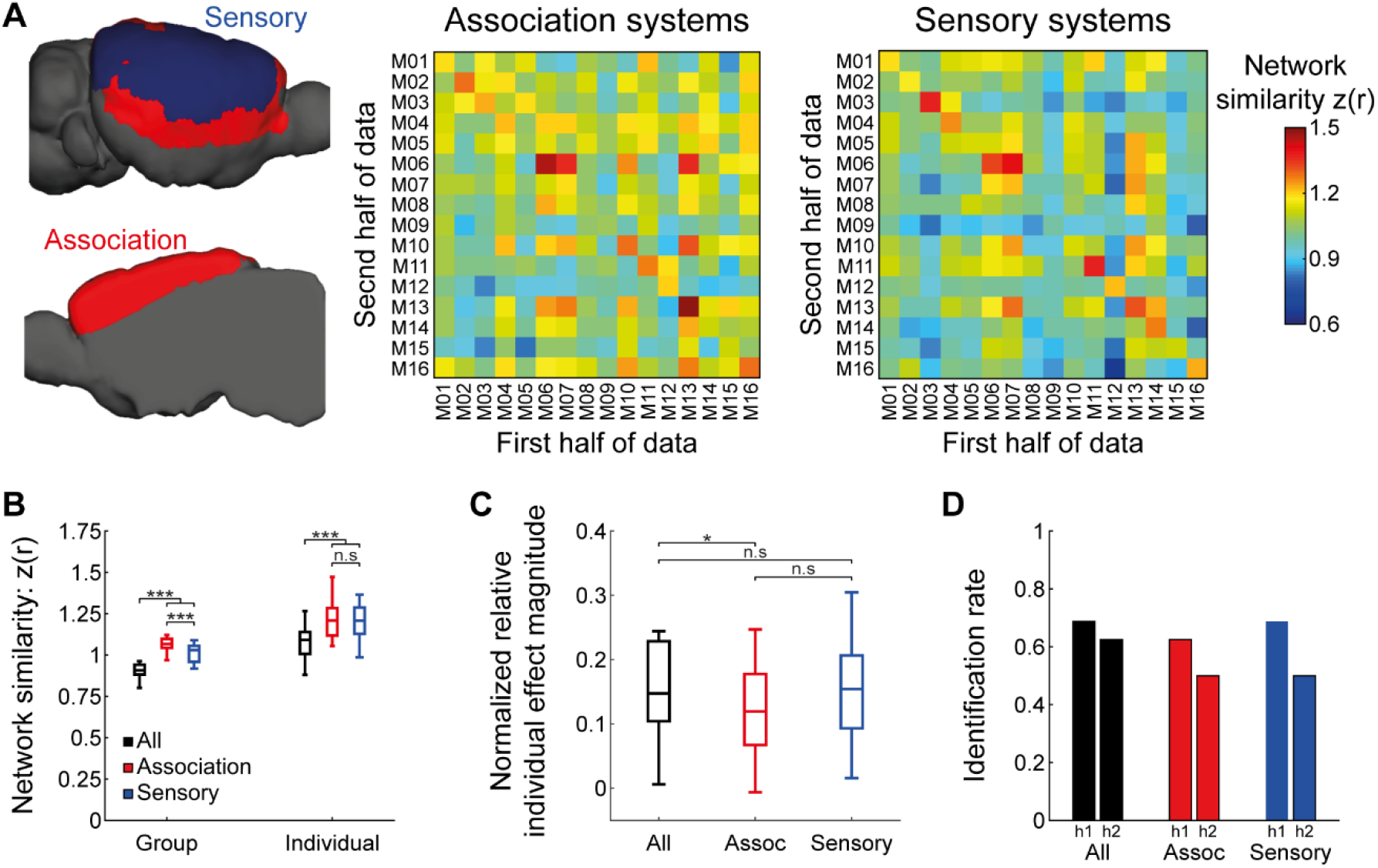
Comparison of individual variation between association and sensory cortical systems. (A) Segmentation of cortical modules to association and sensory networks *(left)* and the similarity matrices between individual connectomes which were limited to either association or sensory regions *(right).* (B) Comparison between group and individual similarities of the full connectome (All) and the connectomes built using only association (Assoc) or sensory regions demonstrates that the full connectome is characterized by lower similarity values at both Group and Individual levels (***P < 0.001). Boxplots represent the median (center line), interquartile range (box limits) and 1.5 × interquartile range (whiskers). (C) Comparison between normalized relative individual effect magnitude of the different connectomes reveals no difference between sensory and association networks. Comparison to the full connectome revealed that it is characterized by a stronger effect of individuality relative to association networks (*P < 0.05). (D) Identification rates of individual mice using the three different connectomes, demonstrate slightly increased performance in the full connectome with minimal differences between association and sensory networks. h1, first half of data; h2, second half.

### Individual brain-behavior relations in mice

After characterizing individual variation in functional connectivity in the mouse cortex, we sought to examine whether it can predict behavioral phenotypes. Therefore, we used connectome-based predictive modelling (CPM)^32^ to link between functional connectivity profiles and behavioral performance in the rotarod task.

Examining rotarod performance, we found prominent variability within the group with different mice presenting wide range of latencies to fall (**Fig. 4A**). This behavioral variability could be predicted by the functional connectivity data using CPM (**Fig. 4B**) as leave-one-out crossvalidation (LOOCV) analysis demonstrated good correspondence (*r*_(16)_ = 0.51) between predicted and observed mean latencies to fall. To formally test the goodness of prediction, we shuffled the behavioral data 1000 times and ran LOOCV analyses on shuffled data to extract significance level (P = 0.021), which confirmed significant prediction. Importantly, latency to fall values were not correlated with individual average head motion estimates during scanning (*r*_(16)_ = −0.27, P = 0.27), confirming that the prediction is not artifactually increased by motion patterns.

**Figure 4:**
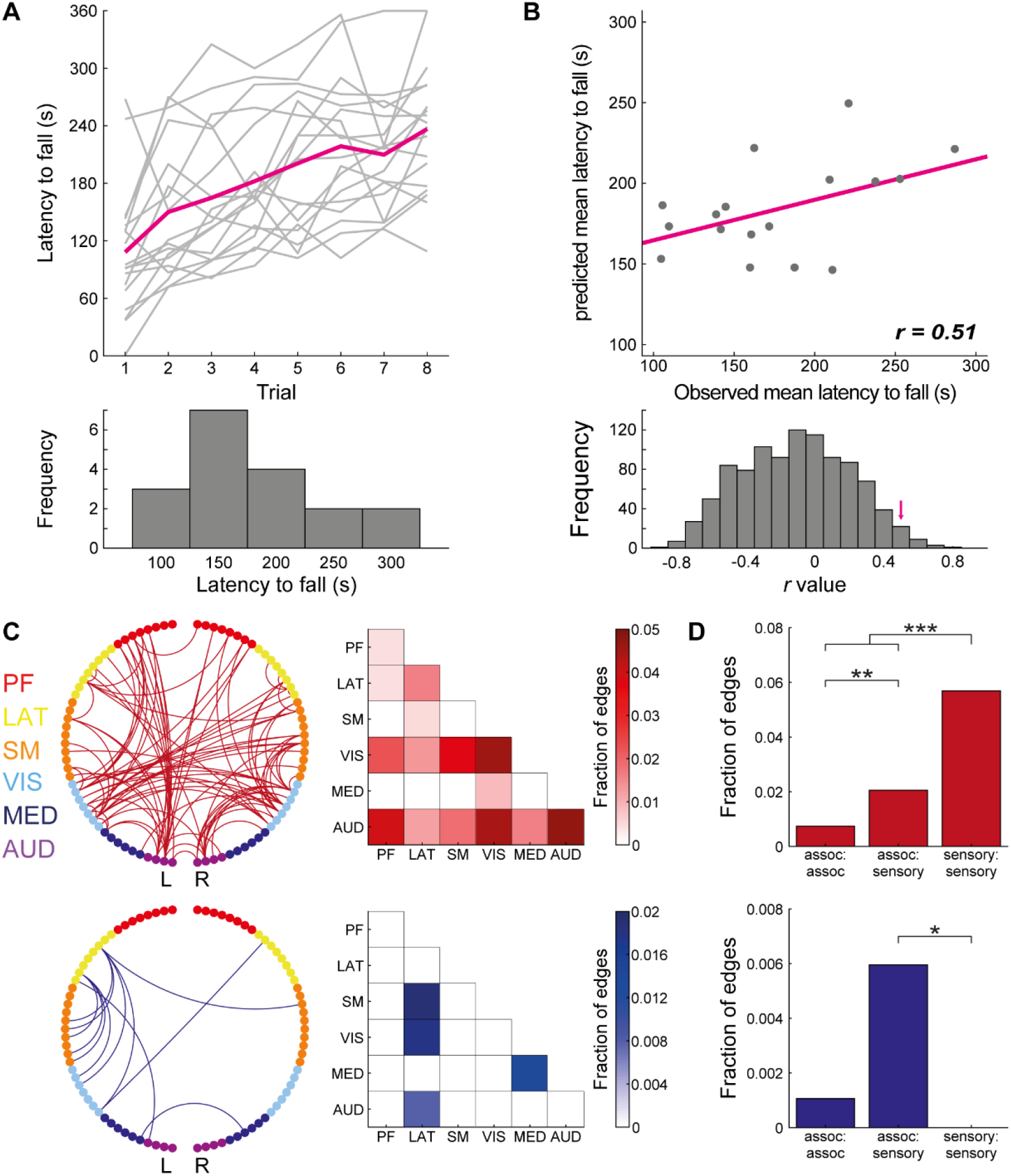
Connectome-based predictive modeling of individual performance in the rotarod task. (A) individual *(light gray)* and group average *(magenta)* performance in the rotarod task *(top),* demonstrate considerable variability in the mean latency to fall of different mice in the group *(bottom).* (B) Relations between observed and predicted mean latency to fall values show close agreement *(top).* A formal statistical analysis was done using a shuffling analysis in which connectome-based predictive modeling was applied to shuffled rotatrod data, the original *r*-value *(magenta arrow’)* designates the 21 highest value out of 1000 iterations. (C) Circle plot *(left)* and matrix *(right)* representations of edges that were positively *(red, top)* or negatively *(blue, bottom)* correlated with rotarod performance in all leave-one-out cross validation iterations. (D) All edges from the previous analysis were divided to three categories, connections between two association regions (assoc:assoc), connections between an association and a sensory region (assoc:sensory) and connections between two sensory regions (sensory:sensory). The results revealed that edges which are positively correlated with the task are biased to sensory:sensory connections *(red,* top), while edges that are negatively correlated with the task are slightly biased to association:sensory connections (*P < 0.05, **P < 0.01, ***P < 0.001). PF, prefrontal; Lat, lateral; SM, somatomotor; VIS, visual; MED, medial; AUD, auditory.

Finally, we explored which functional connections contributed to the model (**Fig. 4C)** and found that positive correlations were more frequent in edges connecting sensory nodes, while negative correlations were less frequent and characterized mainly edges connecting between sensory and lateral association nodes. To formally test this observation we used a set of two-tailed Z-tests for independent proportions (**Fig. 4D**) to test whether significant edges are biased towards connections between association nodes (A:A, n = 946), sensory nodes (S:S, n = 861) or one sensory and one association node (A:S, n = 1848, all comparisons were corrected for multiple comparisons using false-discovery rate). This analysis confirm that positive correlations are more frequent in S:S connections than in A:S (Z = 5, P < 0.001) and A-A (Z = 6.07, P < 0.001) connections, with additional bias towards A:S relative to A:A connections (Z = 2.62, P = 0.009). In contrary, negative correlations were more frequent in A:S connections, demonstrating significant bias relative to S:S connections (Z = 2.27, P = 0.023), and marginally significant bias relative A:A connections (Z = 1.87, P = 0.061); comparison of A:A and S:S connections revealed no difference (Z = 0.95, P = 0.34). Collectively, the data establish that fcMRI can be used to characterize individual brain-behavior relations.

## Discussion

In this study, we characterized individual variation in the functional organization of the mouse cortex. We found evidence for a unique and reliable functional fingerprint, that allows identification of individual mice from a group. Then, we demonstrated that identification rates increase with the amount of data per mouse, indicating the repeated-measurement experimental design is required for precise characterization of mouse-specific connectomes. Comparing individual variation between sensory and association networks, we found that the differences observed between those cortical systems in humans are not well recapitulated in mice. Finally, we show that variance in functional connectivity, especially between sensory cortices, can explain behavioral variability in the rotarod task. Collectively, these findings lay the foundations for studying the functional organization of the mouse brain in health and disease at the level of the individual animal.

While the field of precision fMRI has recently emerged in human brain research^4,7,12,33,34^, rodent data are still analyzed at the group level^35,36^. A major challenge in precision fMRI is the need of large amount of data per subject ranging from 50 to 100 minutes based on the studied brain structure^37^. Such amount of data has yet to be achieved in anesthetized mice in which acquisition is predominantly between several minutes and up to 40 minutes per session^38,39^ and a substantive repeated-measurement design is not feasible due to the difficulties that arise from repeated ventilation and catheterization and the potential impact of prolonged anesthesia. On the other hand, experimental setup for awake fcMRI^27,40–42^ can support such repeated-measurement designs. Therefore, despite the controversy on optimal scanning procedure in rodents and the lack of standardization in the field^43^, awake but not anesthetized fcMRI is necessary for repeated-measurement experimental designs that enable precision fcMRI analysis in individual mice.

Comparison between the findings of this work and the two seminal human studies of Finn^12^ and Gratton^7^ shows that while the central findings of those studies are recapitulated in mice, there are also some important differences. First, identification rates in our mouse cohort are lower and the effect magnitude of individuality is more modest in mice, in which cortical organization is strongly dominated by group shared features. Moreover, the heterogeneity in individual variation among brain networks was not recapitulated in mice, demonstrating similar patterns in sensory and association systems. While these differences might be explained by the complex organization of the human cortex^44^, expansion of association networks in humans^45,46^ or qualitative differences in anatomically homologues high-order structures^27^, we cannot exclude the genetic homogeneity in our cohort reduces the contribution of induvial features, as genetics was previously shown the shape functional connectivity^47^. In addition, some differences are perhaps related to the areal parcellation that was used for defining nodes in the connectome. In humans, connectomes are defined based on functional areal parcellation resulted in several hundred seed regions^48–51^. Since such algorithms have yet to be adapted to mice, and are beyond the scope of the current work, we used anatomical parcellation based on the Allen Mouse Brain Atlas, which is comprised of 86 cortical labels^31,52^. Previous studies in mice showed closed agreement between structural and functional connectivity in the mouse brain^27,53^ including gross cortical organization^54^. Moreover, using the Allen Mouse Brain Atlas we have recently shown that structural connectomes can predict their functional counterpart^29^, suggesting that this anatomical parcellation is functionally relevant.

A major limitation of precision fMRI in humans is the restriction to one level of analysis, as it is limited to large-scale organization^55^. As a result, it is hard to link individual variation in functional connectomes to brain function and dysfunction. In contrast, tools available in rodents support causal manipulation of brain activity using molecular techniques such as chemogenetics or optogenetics, which can be combined with fMRI^39,56–60^. However, heretofore such analyses were restricted to the group level. Future studies can use the individual functional connectome to predict the effect of causal control similar to the way human resting-state fMRI is used to predict individual task fMRI activation pattern^9^, linking the different levels of organization and uncovering sources of individual variation. In addition, rodent models of disease are commonly studied using fcMRI^21,22,61^, which allows direct translation to humans^20,28^, as well as studying the relations between functional connectivity and behavior^23–26^. Such studies can utilize the approach presented here to follow the trajectory of individual animals during development, aging or after treatment, as well as the CPM, which provide a data-driven alternative to predict behavior based on fcMRI data^32^, instead of following a hypothesis-driven behavioral correlations of functional connections between specific seed regions.

Together, our results establish the feasibility of precision fMRI approach in studying the mouse functional connectome, indicating that individual variation in the organization of cortical networks is likely to extend across the mammalian class in general, and is not restricted to primates. Given this foundation, future mouse fMRI studies can follow the human neuroimaging community by moving from group-level inferences to the level of the individual animal. Such transition can be highly beneficial for mechanistic investigation of brain organization, as well as for pre-clinical studies of neuropsychiatric disorders in the context of personalized medicine.

## Data availability

Behavioral and Imaging raw data, as well as custom-written codes used in this study are available in BIDS format on OpenNeuro, https://openneuro.org/datasets/ds002307. Codes for identification and connectome-based predictive modeling adapted from the work of Finn et al.^12^ and Shen et al.^32^ which were previously published: https://www.nitrc.org/projects/bioimagesuite.

## Acknowledgments

This research was supported by Israel Science Foundation (770/17), the Ministry of Science & Technology, Israel & Ministry of Europe and Foreign Affairs (MEAE) and the Ministry of Higher Education, Research and Innovation (MESRI) of France, the Adelis Foundation, and the Prince Center. We thank Technion’s Biological Core Facilities and Edith Suss-Toby for her assistance with MRI, and the Technion Preclinical Research Authority, and Nadav Cohen for assistance with animal care.

## Online methods

### Mice, surgical procedures, behavioral training

All procedures were conducted in accordance with the ethical guidelines of the National Institutes of Health and were approved by the institutional animal care and use committee (IACUC) at Technion. A detailed description of the experimental design was previously published^1^. Briefly, 19 first generation B6129PF/J1 hybrid mice (males, 9–12 weeks old) were implanted with MRI-compatible head-posts and housed in reversed 12 h light/dark cycle. Then, acclimatized to awake fMRI during passive wakefulness^2^ and underwent seven 45 min long awake imaging sessions, followed by one structural MRI session under anesthesia (not used in the current study). A week after functional MRI data were acquired, mice underwent a 2-day rotarod testing.

### Image acquisition

MRI scans were performed at 9.4 Tesla MRI (Bruker BioSpin GmbH, Ettlingen, Germany) using a quadrature 86 mm transmit-only coil and a 20 mm loop receive-only coil (Bruker). Each awake fMRI session started with a brief anesthesia (5% isoflurane) to allow proper mounting to the custom-made cradle^2^. Mice typically were alert within less than a minute. Mice had ~15 min to fully recover from the anesthesia during scanner calibrations and acquisition of a short low-resolution rapid acquisition process with a relaxation enhancement (RARE) T1-weighted structural image (TR = 1500 ms, TE = 8.5 ms, RARE-factor = 4, FA = 180°, 30 coronal slices, 150 × 150 × 450 μm^3^ voxels, no interslice gap, FOV 19.2 × 19.2 mm^2^, matrix size of 128 × 128). Then, four spin echo EPI (SE-EPI) runs were acquired (TR = 2500 ms, TE = 18.398 ms, 200 time points, FA = 90°, 30 coronal slices, 150 × 150 × 450 μm^3^ voxels, no interslice gap, FOV 14.4 × 9.6 mm^2^, matrix size of 96 × 64) before mice were returned to their home cages.

### Rotarod

To assess general motor function, we used the rotarod task^3^ in which mice (up to five at once) walk on a rotating rod (ENV-575MA, Med Associates, St. Albans, VT) while the speed of rotation is accelerating from 16 to 40 rounds per minute during the period of six minutes. Each mouse was trained for two consecutive days over four trials per day with an inter trial interval of 15 min. The latency between the beginning of each trial and falling time was calculated to extract individual learning curves and later averaged across trials to extract a single value for overall task performance per mouse.

### MRI data preprocessing

Functional data were preprocessed as previously described^1,2,4^ including removal of the first two frames for T1-equilibration effects, compensation for slice-dependent time shifts, rigid body motion correction, registration to a downsampled version of the Allen Mouse Brain Atlas (AMBC CCFv3)^5^, and intensity normalization. At this stage, one mouse was excluded due to susceptibility artifacts in the parietal cortex, and additional three sessions were also excluded due to ghosting artifacts in the EPI.

After this general fMRI preprocessing, an fcMRI-specific preprocessing was performed. First, data underwent motion scrubbing to remove motion-related artifacts^6^. Censoring criteria were frame displacement of 50 μm and temporal derivative root mean square variance over voxels of 150% inter-quartile range above the 75^th^ percentile, with an augmented mask of one additional frame after each detected movement and censoring of sequences with less than five included frames. Runs with less than 50 frames and sessions with less than 192 frames (8 min) were excluded (a total of six sessions). The average number of included sessions per mouse was 6.33 ± 0.84 (mean ± SD) and the average total included time per session was 19.31 ± 3.67 min per session. After motion scrubbing, preprocessing continued with demeaning and detrending, nuisance regression of six motion parameters, ventricular and white matter signals, and their first derivatives, temporal filter (0.009 < f < 0.08 Hz), and spatial smoothing (Gaussian kernel with FWHM of 450 μm).

### Construction of the functional connectome

To construct the functional connectome of each mouse, we used the AMBC Atlas to define 86 regions in the mouse cortex (43 per hemisphere), which were classified into six different modules (Prefrontal, Lateral, Somatomotor, Visual, Medial and Auditory) based on their anatomical connectivity patterns^5^. Labels were registered to the native fMRI resolution using the nearest neighbor interpolation^7,8^. The very deep and superficial aspects of each label were removed to minimize partial-volume effects. In addition, posterior parts of the retrosplenial and primary visual cortices were also removed due to inconsistent registration in these areas.

After defining connectome nodes, we extracted their time courses in each session of each mouse, and calculated the Fisher’s Z transformed Pearson correlation (*r*) values^9^, resulting in a 86 × 86 connectivity matrix per session. Then, these matrices were split to two halves of three sessions per mouse for similarity and identification analyses (minimizing the difference in number of included frames per half) or averaged across all sessions of each mouse for the connectome-based predictive modelling (CPM) analysis. Mice with less than six valid sessions (n = 2) were included only in the CPM analysis. In mice with seven valid sessions (n = 9), the session with the highest movement was excluded from the similarity and identification analyses in order to match the amount of data per half.

### Similarity and identification analyses

To quantify individual variation and perform connectome-based fingerprinting we adapted algorithms developed in humans to estimate network similarity^10^ and identification rates^11^. Both procedures are based on the construction of a similarity matrix, in which each cell is the Fisher’s Z transformed correlation between val7ues in all 3655 edges in two connectomes, columns represent the first half of data and rows represent the second half of data. For quantification of network similarity, values along the diagonal represent individual similarity, which is the correlation between the connectomes built for the same mouse from the halves of data. Group similarity is defined as the average of values in a combined row and column vector (excluding the value along the diagonal). Group and individual similarity values were compared using paired student *t*-test or repeated-measures ANOVA after normality was tested using the Lilliefors test.

The identification procedure is a more stringent analysis for fingerprinting and quantifies the fraction of mice in which the individual similarity is higher than any other value in their row or column. To formally test whether the observed identification rates are significant, we compared them to the distribution of rates in 1000 iterations in a shuffling analysis, in which we assigned random identity to connectomes of one half of the data, and examined whether this random identity can be detected using data from the other half of data. To test whether individual variation differs between association and sensory networks in the mouse cortex, we constructed connectomes limited to either association (n = 44) or sensory (n = 42) modules. Then, we ran the similarity and identification analyses as described.

### Quantifying the amount of data needed to study individual variation in mice

To characterize the amount of data needed for studying individual variation in mice, we examined the effects of the number of sessions included in the construction of each connectome on network similarity and identification rate. We built two connectomes per mouse using one, two or three sessions per connectome, and examined all possible combinations. Then, we replicated the similarity and identification analyses as described.

### Quantifying edgewise contributions to identification

Edge-based analyses examine which connections contribute more to successful identification. A detailed description of the calculation of group consistency (Φ) and differential power (DP) was previously published^11^. Briefly, each connectome underwent *z*-normalization, and the edgewise product was calculated for each edge in all pairs of connectomes. Group consistency values are the average edgewise products from the comparisons of the two connectomes of each mouse. In contrast, DP represents the empirical probability that edgewise product of two connectomes from the same mouse is higher than the edgewise product of two connectomes from different mice. The top 1% of DP and Φ were presented in either circle plots or matrix plots grouped to modules ^12^.

### Connectome-based predictive modelling (CPM)

To link individual variation in the functional connectome to behavioral variability in the rotarod task, we followed a previously published detailed protocol^12^. We used a leave-one-out crossvalidation approach in which Spearman correlation was calculated between functional connectivity in each edge in the connectome and rotarod mean latency to fall values. Then, significant (P < 0.05) positive and negative correlations were selected, summarized to two values per mouse, which were combined in a single regression model. This model was used for prediction of rotarod performance in the one testing mouse in each one of the 18 iterations. Finally, the correlation between predicted and observed behavioral measures was calculated and formally tested by comparing it to the distribution of correlation in 1000 iterations in which rotarod performance was randomly assigned to mice^12^. To characterize the edges that contributed to the prediction, we extracted the edges that were included in the model in all iterations and presented them in circle and matrix plots. A set of Z-tests for independent proportions was used to examine whether the contributing edges were biased to connections between association or sensory regions.

